# Landscape heterogeneity as a main driver of avian population dynamics

**DOI:** 10.64898/2026.05.19.726359

**Authors:** Katarzyna Malinowska, Tomasz Chodkiewicz, Lechosław Kuczyński

## Abstract

The ongoing decline in biodiversity highlights the need for understanding the causes of population changes. This study uses 25-year, large-scale monitoring dataset to investigate the influence of climate and landscape structure on the annual population growth rates of 84 bird species across Poland. Our methodological framework involves the spatiotemporal decomposition of these environmental drivers to decouple demographic effects of long-term carrying capacities from the short-term effects of environmental perturbations. Using species-specific demographic models followed by a community-wide meta-analysis, we evaluated how individual species responses scale up to shape community-level dynamics. The results reveal significant variation in species-specific responses to individual drivers. At the community level, our findings suggest that bird populations are mainly regulated by the long-term spatial constraints rather than short-term disturbances. Persistent environmental heterogeneity had the strongest positive demographic effect on birds, followed by temperature, forest dominance over croplands, and precipitation. In contrast, rapid temporal shifts in environmental heterogeneity and precipitation anomalies negatively affected population growth, whereas urbanisation consistently exerted a negative effect across both spatiotemporal dimensions. Our results highlight the significance of protecting existing heterogeneous and ecotonal habitats, as well as the need to incorporate features that enhance habitat heterogeneity into urban development.

**Article impact statement:** Preserving heterogeneous habitats is essential for the conservation of bird populations.

## Introduction

Biodiversity is declining across numerous taxonomic groups, affecting both terrestrial and marine organisms worldwide (Ceballos et al., 2017; McCauley et al., 2015). Among vertebrates, this crisis has been particularly well documented in birds, perhaps due to their wide distribution and ecological diversity, which makes them valuable indicators for assessing environmental quality (Fraixedas et al., 2020).

Over the past several decades, most farmland birds have experienced dramatic long-term population declines, which at the continental scale is an undeniable loss of biodiversity (Burns et al., 2021; Gregory et al., 2005, 2023). On the other hand, many bird species associated with woodland and forest habitats, have maintained stable or increasing populations at the continental level (Gregory et al., 2023; Reif, 2013), but at finer scales these patterns show substantial variability (Ram et al., 2017; Rigal et al., 2023). Other functional groups, such as urban species, show a highly region-dependent trends (Reif, 2013; Rigal et al., 2023).

Identifying macroecological drivers underlying these both positive and negative population trends is an important step before implementing effective conservation strategies and recovery plans. Changes in landscape structure are among primary factors affecting bird population dynamics. Decline in environmental heterogeneity driven by land-use change or management practices can contribute to significant biodiversity loss (Benton et al., 2003; Fahrig et al., 2011). It often resulted from agricultural intensification, which is recognised as one of the main causes of bird population declines, deteriorating habitat quality in multiple ways (Guerrero et al., 2024; Rigal et al., 2023). In particular, rapid transitions from extensive to intensive agriculture in many Central and Eastern European (CEE) countries led to the drainage of meadows and peat bogs, the removal of hedgerows, shrubs, and other non-cultivated areas, which reduce the amount of ecotonal habitats and promote large-scale monocultures (Emmerson et al., 2016). Another significant landscape change observed in most CEE countries is the expansion of forest cover, largely driven by large-scale agricultural land abandonment following the post-communist transition (Janus & Bozek, 2019). This process may temporally benefit bird communities by increasing landscape complexity and creating novel habitats (Tryjanowski et al., 2011). Consequently, secondary natural succession and forest expansion resulting from land abandonment contribute to significant turnover in bird species composition (Broughton et al., 2022; Corkery et al., 2020). Finally, the accelerating pace of urbanisation and increasing anthropogenic pressure have a negative impact on avian populations, often leading to the destruction and fragmentation of natural habitats (Rigal et al., 2023). Processes such as urban sprawl imply a negative effect on species diversity and lead to the homogenisation of species communities (McKinney, 2006; Sidemo-Holm et al., 2022). Another type of environmental factor shaping bird population dynamics is climate change. Its effect on bird populations is not uniform, as increasing temperatures often favour thermophilic species, while populations of cold-dwellers tend to decline, which create distinct groups of ecological ‘winners’ and ‘losers’ (Davey et al., 2012; Pearce-Higgins et al., 2015; Stephens et al., 2016).

The drivers described above can affect bird communities through two distinct pathways: as long-term processes that shape the baseline characteristics of the landscape (i.e. the environmental carrying capacity), and as short-term disturbances that cause temporary deviations from these baseline conditions (Begon et al., 2005). Distinguishing between these two mechanisms provides a clearer identification of the underlying causes of population changes and helps determine where conservation efforts should be focused. Consistently, decomposition of environmental factors into long-term spatial means and temporal anomalies has been applied in similar avian studies (Bradter et al., 2022; Jørgensen et al., 2016).

Understanding the causes of bird population changes across Europe requires consideration of country-specific socio-ecological context. This is particularly relevant for Central and Eastern Europe, where political and economic transitions have led to profound differences in agricultural practices, land use, and urban development compared to Western Europe (Kuemmerle et al., 2016; Tryjanowski et al., 2011). Considering a similar historical background, findings of our study may offer valuable insights relevant to other post-communist countries.

The study evaluates the impact of environmental drivers on annual avian population growth rates using a large-scale, 25-year monitoring dataset encompassing 84 bird species across Poland. Our analytical framework focuses on climatic factors, such as temperature and precipitation, and dominant landscape structure in Poland, which capture forest – cropland gradient, environmental heterogeneity, and urbanisation. Each driver was decomposed into a long-term spatial baseline and a short-term temporal disturbance, to decouple their distinct impact on population dynamics. We employed demographic models to quantify species-specific responses, followed by a meta-analysis of the resulting effect sizes to assess the community-wide effect of environmental drivers. In this study we aim to identify environmental factors influencing bird growth rates and to explain mechanisms underlying these effects, specifically whether they shape a long-term carrying capacity or act as short-term environmental perturbations. Moreover, we aim to investigate how species-specific demographic responses scale up to determine the overall stability and composition of the community. Ultimately, we intend to provide practical recommendations for habitat management and biodiversity conservation.

## Materials and Methods

### Bird population data

In the study, we used bird count data from the Common Breeding Birds Survey (polish acronym MPPL), covering the whole Poland. Data from the years 2007–2024 were obtained from the State Environmental Monitoring coordinated by the Chief Inspectorate of Environmental Protection (GIOŚ), while data from the earlier period (2000–2006) were obtained from the Polish Society for the Protection of Birds (OTOP), providing a continuous 25-year time series of bird counts. MPPL is also part of the Pan-European Common Bird Monitoring Scheme (PECBMS, Brlík et al., 2021), and its methodology follows a well-established protocol: bird counts are conducted twice a year during the breeding season on 1×1 km plots selected in stratified random sampling. On each of these plots, ornithologists follow two separate transects and records all seen or heard adult birds. As a measure of relative species population size for each plot, we used the higher of the two counts obtained during the early and late breeding-season visits.

The MPPL provides annual information on approximately 110 common bird species in Poland. However, for the purpose of the analysis and credibility of results we used a subset of these species, and we excluded strictly colonial species (such as gulls, terns, and the rook), waterbirds and birds of prey. Our aim was to focus the analysis on species that can be recorded in multiple territories within a 1 km² plot and whose distribution across the landscape is spatially dispersed rather than aggregated by colonies or restricted to water bodies. We have excluded all birds of prey, because their home ranges often extend far beyond the 1 km² monitoring survey plots, making it difficult to reliably assign individuals to specific sampling units and potentially biasing estimates of species occurrence. Eventually, the set of 84 most common bird species occurring in Poland was used in the analysis, with majority of them being passerine birds (for a detailed list of species see Table S1).

### Environmental covariates

Full details regarding data acquisition and spatial processing of environmental covariates are provided in the Supplementary Material.

#### Land-cover data

Annual land cover data (2001–2024) were obtained from the 500-m resolution MODIS Land Cover Type product (MCD12Q1 v6.1; Friedl & Sulla-Menashe, 2022), utilising the 17-class IGBP classification scheme (Sulla-Menashe & Friedl, 2022). Data were acquired via NASA AppEEARS service, then mosaicked and cropped to the area of Poland using the *terra* package in R (Hijmans et al., 2026). To quantify fractional cover, binary mask for each IGBP class was reprojected to our 1-km analytical reference grid using bilinear interpolation. That way we obtained continuous proportional area values for seven land cover classes, calculated for each grid cell.

#### Climatic data

Bioclimatic data originate from TerraClimate, which provides a monthly climate information with a spatial resolution of ∼4 km (Abatzoglou et al., 2018). The data were imported into R and reprojected at a spatial resolution of 1 km using bilinear interpolation and functions from the *terra* package (Hijmans et al., 2026). For this study, we used information on the average annual temperatures and log-transformed total precipitation for each year between 2001 and 2024.

#### Landscape dimensionality reduction

To address the multicollinearity inherent in spatiotemporally autocorrelated landscape features, we applied sparse principal component analysis (SPCA) to the seven satellite-derived land-cover classes using the *elasticnet* package in R (Zou et al., 2006). As all land-cover variables were measured on an identical proportional scale (0–100%), the SPCA was computed on the covariance matrix rather than the correlation matrix, which preserved the true magnitude of variance inherent to each class. Principal components were calculated using a sparsity penalty (L1 regularisation) to set weak loadings exactly to zero. The resulting axes are therefore not only mathematically independent, but also biologically interpretable.

We extracted three primary sparse components, which together explained 86.6% of the spatiotemporal variance in the land cover data (Table 1). The first component (PC1; explaining 63.8% of the variance) was termed the “forest-cropland” gradient. It captured the predominant landscape feature in Eastern Europe, reflecting the transition between arable land (with negative loadings) and dense, uniform forests (with positive loadings). The second component (PC2), which explained an additional 18.7% of the variance, represented the “environmental heterogeneity” gradient and captured the spectrum of landscape diversity. It contrasted uniform monocultures (with negative loadings for structurally homogeneous forests and intensive arable land) with spatially diverse ecotones (with positive loadings for transitional woodlands, open forests and grasslands). Finally, the sparsity penalty isolated urban cover onto an independent third component (PC3), which explained an additional 4.1% of the variance and was labelled the “urbanisation” gradient. It accounted for built-up infrastructure across both rural and urban settlements. The spatial and temporal characteristics of the resulted environmental variables, which were used in further analyses (PC1, PC2, PC3, temperature, and precipitation) are detailed in Table S2.

**Table 1.**
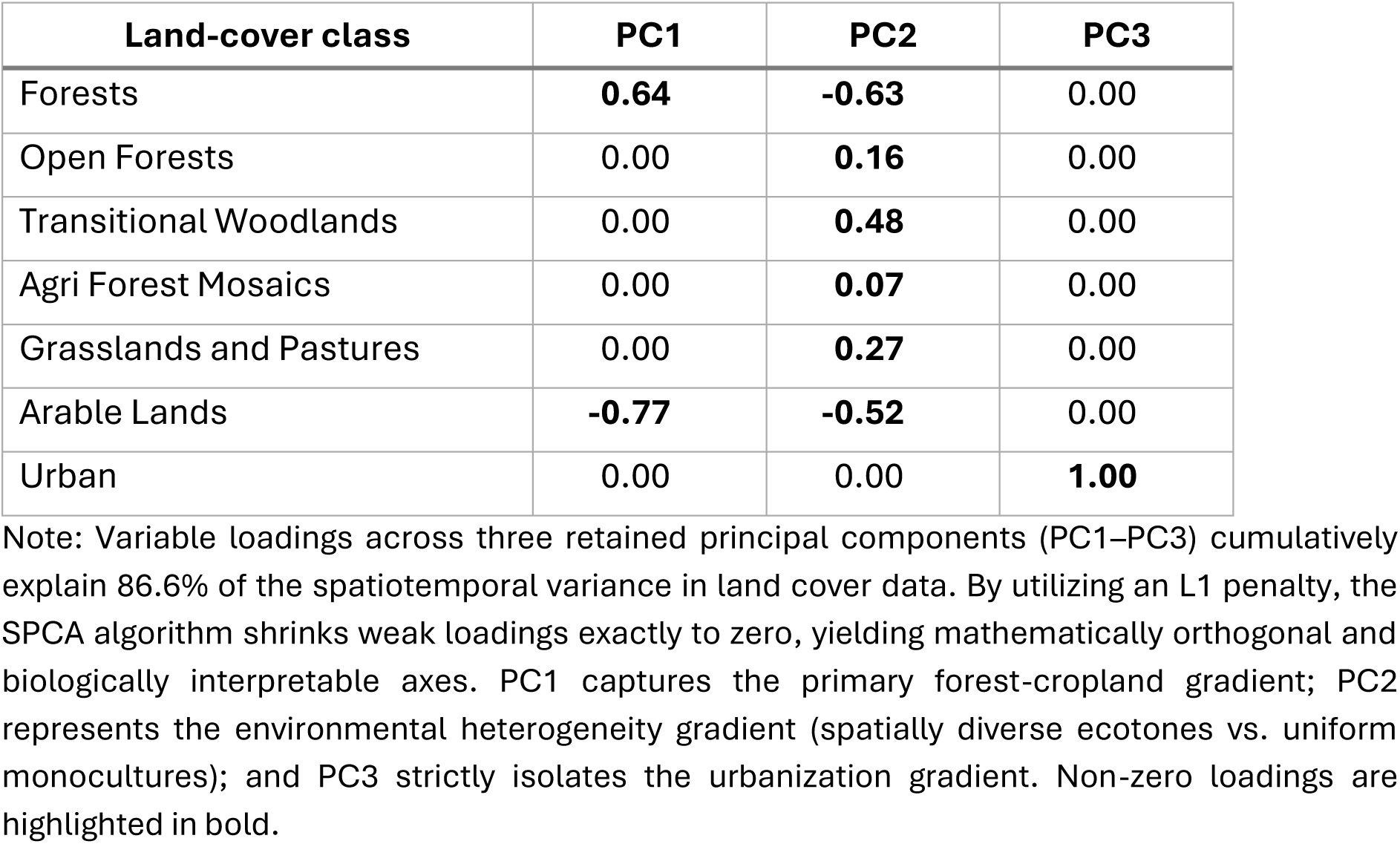
Sparse principal component analysis (SPCA) loadings for satellite-derived landscape variables. Note: Variable loadings across three retained principal components (PC1–PC3) cumulatively explain 86.6% of the spatiotemporal variance in land cover data. By utilizing an L1 penalty, the SPCA algorithm shrinks weak loadings exactly to zero, yielding mathematically orthogonal and biologically interpretable axes. PC1 captures the primary forest-cropland gradient; PC2 represents the environmental heterogeneity gradient (spatially diverse ecotones vs. uniform monocultures); and PC3 strictly isolates the urbanization gradient. Non-zero loadings are highlighted in bold.

| Land-cover class | PC1 | PC2 | PC3 |
| --- | --- | --- | --- |
| Forests | **0.64** | **-0.63** | 0.00 |
| Open Forests | 0.00 | **0.16** | 0.00 |
| Transitional Woodlands | 0.00 | **0.48** | 0.00 |
| Agri Forest Mosaics | 0.00 | **0.07** | 0.00 |
| Grasslands and Pastures | 0.00 | **0.27** | 0.00 |
| Arable Lands | **-0.77** | **-0.52** | 0.00 |
| Urban | 0.00 | 0.00 | **1.00** |

### Spatiotemporal decomposition of environmental drivers

#### Spatiotemporal decomposition of climatic stressors

In contrast to land-cover classifications, which are prone to abrupt, stepwise transitions, climatic variables act as continuous drivers characterised by pronounced interannual variability superimposed on long-term directional trends. To decouple the effects of fundamental climatic niches from these temporal weather dynamics, we applied a standard variance decomposition (Mundlak, 1978) to the annual climatic time series at each site. For both temperature and precipitation, we partitioned the raw annual data into two distinct demographic drivers. Firstly, we calculated the spatial mean (the 24-year average for each site), which defines the long-term climatic carrying capacity. Secondly, we calculated the temporal anomaly for each site by subtracting its spatial mean from its observed annual value. This isolates the temporal variance within a site, representing abnormal weather conditions or resource pulses (e.g., an unusually severe drought or an unseasonably warm spring) that occur in a given year.

#### Spatiotemporal decomposition of landscape trajectories

Satellite-derived land-cover classifications are inherently unstable and often switch between similar classes from one year to the next (Friedl et al., 2010). This “flicker” effect occurs because ecotones and heterogeneous landscapes contain a mixture of cover types within a single satellite pixel, resulting in spectral signatures that are difficult to distinguish. Furthermore, transient natural variability, such as shifting phenology or temporary disturbances (e.g., drought or insect outbreaks), introduces noise into the classification system. To eliminate these artefacts, we applied a trajectory smoothing and signal extraction protocol to the three independent principal components (PC1–PC3) at each site. We modelled the temporal trajectory of each landscape component using generalised additive models (GAMs) via the *mgcv* package in R (Wood, 2017) and parameterised them using penalised B-splines (P-splines).

Land-cover change can manifest as either abrupt anthropogenic conversions, such as clear-cutting and urbanization, or gradual natural succession, such as shrub encroachment. In order to capture both these stepwise transitions and gradual shifts while strictly ignoring interannual classification noise, we applied a first-order penalty (m = 1) to the spline, which shrinks the first derivative towards zero. We coupled it with a null-space penalty (select = TRUE) estimated via restricted maximum likelihood (REML) to ensure that genuinely stable landscapes were constrained to a flat line.

From the resulting noise-free trajectories, we extracted two demographic drivers for each principal component. Firstly, since the temporal predictor (year) had been median-centred prior to smoothing, we extracted the GAM intercept, representing the baseline environmental state at the exact temporal midpoint of the study period (hereafter referred to as the “spatial baseline”). Secondly, we calculated the first derivative (year-on-year difference) of the smoothed trajectory to capture the exact velocity of the environmental change. We termed this dynamic metric the “temporal delta.”

#### Driver standardisation

Following the spatiotemporal decomposition, each of the three landscape features was partitioned into a spatial baseline and a temporal delta. Similarly, the two climatic variables were partitioned into spatial means and a temporal anomalies. These 10 main drivers were then incorporated into demographic models. To enable direct comparisons of effect sizes across different units of measurement, all of the extracted spatiotemporal drivers were standardised independently to a mean of zero and a standard deviation of one (Z-scores). They were then integrated into species-specific demographic models.

### Demographic modelling

#### Model specification and density dependence

Although our theoretical framework focuses on the realised annual population growth rate (*r*) as the demographic response, calculating growth rates in advance using differenced counts can increase observation error and make exact zeros more difficult to handle. Instead, we modelled the raw annual counts (*N*_*t*_) directly using a Tweedie distribution with logarithmic link function, which efficiently handles the zero-inflation and overdispersion that are common in survey data. To convert the measure of abundance into a growth rate, we used the log-transformed count from the previous year at the same site, *ln*(*N*_*t*−1_), as the model offset. As the link function is logarithmic, it effectively shifts the lagged term to the left-hand side of the regression equation (i.e., *ln*(*N*_*t*_) − *ln*(*N*_*t*−1_)). Therefore the model explicitly evaluates the growth rate (*r*). To account for internal population regulation, we simultaneously included the lagged log-abundance as a penalised non-linear smooth term. This flexible Gompertz formulation can potentially accommodate complex demographic realities, capturing both negative density dependence when populations approach local carrying capacity, and positive density dependence (Allee effects) at low abundances. The dual specification ensures that the estimated effects of environmental drivers are separated from the species’ internal population dynamics.

#### Spatiotemporal autocorrelation and detection bias

As observational count data are prone to unmeasured environmental variance, sampling heterogeneity, and imperfect detection, we implemented a rigorous, multi-layered error structure. We included independent random intercepts for survey site and year to capture spatial and temporal heterogeneity. To control explicitly for differences in detection probability and skill level among field ornithologists, we included a random intercept for observer identity. Finally, to account for broad-scale biogeographic gradients and long-term population trends that are independent of our focal landscape and climatic drivers, we fitted two Gaussian process smoothers: a two-dimensional spatial surface for the site coordinates, and a continuous global temporal smoother for the study year.

#### Driver regularisation

To estimate the effects of ten standardised environmental drivers (four climatic and six land-use variables), we entered them into the GAMMs as parametric fixed effects. To prevent residual multicollinearity among these drivers, we applied an L2 (ridge) penalty to the fixed-effects matrix. This regularisation algorithm iteratively shrinks the coefficients of weakly informative drivers towards zero, ensuring that the final extracted effect sizes represent highly stable demographic responses.

#### Ecological interpretation of fixed coefficients

Our models define the realised population growth rate (*r*) as the response variable and utilise standardised predictors (Z-scores), allowing the resulting fixed-effect coefficients to serve as directly comparable measures of demographic effect size. Mathematically, a given coefficient (*β*) represents the slope, i.e. the additive change in a species’ annual growth rate caused by a one-standard-deviation increase in the environmental predictor. Ecologically, however, these coefficients must be interpreted through the dual lens of our spatiotemporal decomposition.

The spatial components of our decomposition represent long-term averages, and therefore slopes against them quantitatively assess the demographic impact of an environment’s carrying capacity. A positive spatial coefficient for a given variable indicates that a species consistently achieves higher realised growth rates in areas where that environmental variable is consistently high. Conversely, temporal components represent short-term local changes, such as landscape transitions or weather anomalies. Consequently, slopes against these temporal variables reflect the demographic response to environmental perturbations. A negative temporal coefficient indicates that a shift in local conditions, such as clear-cutting or an extreme thermal event, acts as a demographic stressor, reducing the population growth rate in that specific year.

Ultimately, as all environmental drivers were standardised to a common scale and regularised via an L2 penalty to prevent variance inflation among correlated predictors, these coefficients represent stable and directly comparable effect sizes. This robust mathematical framework enables us to directly compare the magnitude of long-term habitat constraints with the severity of short-term environmental disturbances.

### Meta analysis

For each species model, we extracted the effect size estimates (regularised *β* regression coefficients) alongside their corresponding variance–covariance matrices. These matrices were then combined into a single block-diagonal matrix. In the next step, we conducted a meta-analysis on the obtained *β* coefficients from all models using *rma.mv* function from the *metafor* package (Viechtbauer, 2010). To account for both the estimation uncertainty and the non-independent covariance structure among the drivers within a given species, we weighted the meta-analysis using the constructed block-diagonal matrix. Environmental drivers were treated as fixed effect moderators, whereas species identity was modelled as a random intercept to account for dependence among effect sizes. The model was fitted using restricted maximum likelihood (REML) estimation, allowing for the calculation of an overall mean effect size and corresponding confidence intervals for each environmental driver across the community.

The analyses were performed in R 4.5.1 software (R Core Team, 2025). The full R code and data used in the meta-analysis part are available at: https://github.com/popecol/Drivers.

## Results

### Species-specific demographic responses

We successfully fitted species-specific demographic generalised additive mixed models (GAMMs) for 84 bird species, encompassing over one million local population transitions (i.e., year-to-year changes in abundance at individual survey sites; median n = 12,270 observations per species). The models demonstrated robust predictive performance, yielding a median adjusted R^2^ of 0.62 (range: 0.23–0.90). Importantly, the non-linear formulation for lagged abundance was strongly supported by the data, with an average effective degree of freedom (EDF) of 2.35 for the density-dependent smooth term across the modelled species. Since an EDF of exactly 1.0 indicates a purely linear relationship, the high mean EDF reveals widespread non-linear population regulation within the community, a pattern that would otherwise be obscured by strict linear Gompertz assumptions.

Evaluation of the individual fixed-effect coefficients (*β*) revealed substantial species-level heterogeneity (Figs. S1–S10, Supplementary Material), confirming that distinct life histories and ecological niches mediate environmental responses. The magnitude of these coefficients reflects the extent of species-specific specialisation. Those with high absolute *β* values demonstrate strict environmental specialisation and occupy narrow ecological niches. Conversely, near-zero coefficients indicate ecological generalism, reflecting species that occupy broad niches and tolerate wide environmental gradients.

Spatial means, reflecting local carrying capacity, exhibited high demographic divergence. For example, the long-term thermal niche generally had a positive effect on the community, however individual species responses ranged from strongly negative (β=−0.70, indicating strict specialisation for cooler microclimates) to highly positive (β=1.04, indicating strong thermophilous specialisation) (Fig. S9). Similarly, the spatial baseline of the forest-cropland gradient produced polarised demographic responses, with slopes ranging from −1.85 to 1.66, reflecting a spectrum from cropland specialists to strict forest specialists (Fig. S1).

Responses to short-term environmental perturbations were smaller in magnitude, yet equally heterogeneous in direction. For instance, annual temperature fluctuations resulted in a temporary increase in the population size for some species (β=0.14), while leading to a sharp decline for others (β=−0.14) (Fig. S10). This bidirectional variance in temporal effect sizes reflects differences among species in their ability to buffer against short-term environmental stressors.

### Community-wide demographic responses

The meta-analysis revealed that long-term spatial constraints had a much stronger demographic impact on the dynamics of regional bird communities than short-term environmental disturbances (Table 2, Fig. 1).

**Figure 1.**
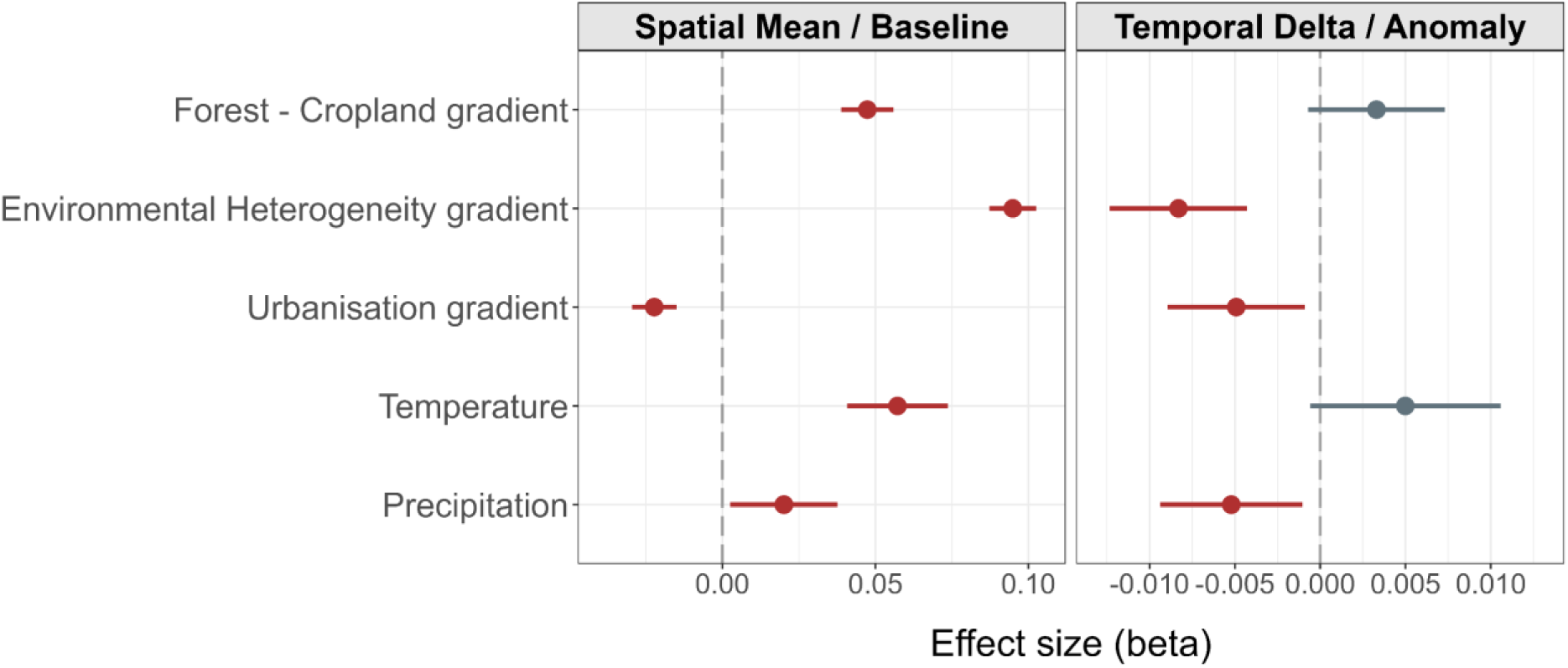
Meta-analytic mean effect sizes (*β*) of standardised environmental drivers on the annual population growth rates of 84 bird species. Results are partitioned into spatial baselines (representing long-term carrying capacity constraints) and temporal anomalies (representing short-term environmental perturbations). Points represent the estimated mean coefficients, and horizontal bars indicate 95% confidence intervals. Statistically significant effects (*p* < 0.05) are highlighted in red, whereas non-significant effects are shown in grey.

**Table 2.** Results of the meta-analysis estimating mean effect sizes (*β*) for the impact of standardised environmental drivers on annual population growth rates across 84 bird species. Note: The meta-analytic model includes species as a random intercept, and utilises a block-diagonal variance-covariance matrix to account for non-independence among driver estimates within species. Statistically significant results (*p* < 0.05) are shown in bold.

| Driver name | Mean beta | p-value | Lower 95% confidence interval | Upper 95% confidence interval |
| --- | --- | --- | --- | --- |
| **Spatial Baseline / Mean** |  |  |  |  |
| Forest – Cropland gradient | 0.047 | **<0.0001** | 0.039 | 0.056 |
| Environmental Heterogeneity gradient | 0.095 | **<0.0001** | 0.087 | 0.103 |
| Urbanisation gradient | -0.022 | **<0.0001** | -0.030 | -0.015 |
| Precipitation | 0.020 | **0.025** | 0.003 | 0.038 |
| Temperature | 0.057 | **<0.0001** | 0.041 | 0.074 |
| **Temporal Delta / Anomaly** |  |  |  |  |
| Forest – Cropland gradient | 0.003 | 0.106 | -0.001 | 0.007 |
| Environmental Heterogeneity gradient | -0.008 | **<0.0001** | -0.012 | -0.004 |
| Urbanisation gradient | -0.005 | **0.017** | -0.009 | -0.001 |
| Precipitation | -0.005 | **0.015** | -0.009 | -0.001 |
| Temperature | 0.005 | 0.080 | -0.001 | 0.011 |

Among the spatial baselines, long-term landscape composition had the strongest influence on population growth rates. Of the five tested effects, four were positive and one was negative. The strongest positive effect was habitat heterogeneity (β = 0.095, SE = 0.004, p < 0.0001), followed by the spatial mean of temperature (β = 0.057, SE = 0.008, p < 0.0001), forests dominance over croplands (β = 0.047, SE = 0.004, p < 0.0001), and the spatial mean of precipitation (β = 0.020, p = 0.025). Urbanisation was the only variable with a negative effect on regional growth rates (β = −0.022, SE = 0.004, p < 0.0001). Because individual species demonstrated varying degrees of specialisation, these strong, uniform shifts at the meta-analytic level indicate that persistent environmental conditions actively shape the regional community. Specifically, a high proportion of heterogeneous (ecotonal) habitats, the dominance of forests over croplands, and persistently warmer or wetter conditions favour population growth, while highly urbanised or cropland-dominated areas actively constrain it.

Although the magnitude of demographic responses to short-term local environmental perturbations was markedly lower than that of spatial constraints, they revealed critical community-wide vulnerabilities. Of the five tested effects, three were significantly negative, while two lacked a uniform direction across the community. Among the significant effects, rapid shifts in habitat heterogeneity had the strongest negative influence, indicating a sudden, presumably anthropogenic disturbances (β=−0.008, SE = 0.002, p < 0.0001). It was followed by increased urbanisation (β=−0.005, SE = 0.002, p = 0.017) and precipitation anomalies, i.e., exceptionally wet years (β=−0.005, SE = 0.002, p = 0.014), both actively depressed annual growth rates across the communities. In contrast, transitions in the forest-cropland gradient (β=0.003, p = 0.106) and temperature anomalies (β=0.005, p = 0.080) lacked a uniform direction.

## Discussion

Our findings demonstrate that avian population dynamics are predominantly regulated by long-term spatial constraints rather than short-term environmental disturbances. By decomposing complex environmental drivers into static spatial baselines (representing carrying capacities) and temporal deviations (reflecting habitat transitions and annual climatic anomalies), we identified a dual spatiotemporal mechanism shaping the regional bird community. At the species level, demographic responses to temporal disturbances are highly heterogeneous, reflecting the distinct ecological niches of different species. At the community level, however, these diverse short-term responses often cancel each other out, indicating that the overall community composition is filtered by spatially persistent environmental states, rather than by temporal environmental changes. Specifically, the carrying capacities of the studied avian community are primarily increased by landscape heterogeneity, forest dominance, and climatic suitability (i.e., warmer and wetter conditions), while being severely limited by high urbanisation. Together, these spatial drivers act as fundamental macroecological filters that strictly dictate community composition.

### The dominance of long-term spatial constraints

Our results indicate that within the landscape domain, habitat heterogeneity and the dominance of forest over croplands are the strongest spatial drivers of avian population growth. Highly heterogeneous and ecotonal landscapes provide a wide variety of ecological niches by offering diverse nesting sites, foraging opportunities, and refuge (Fahrig et al., 2011; Tews et al., 2004). Additionally, through landscape complementation (Dunning et al., 1992) more complex habitats can accommodate a larger number of specialist species, which increases the overall carrying capacity. Similarly, the positive demographic response to forest dominance reflects the high structural complexity (Betts et al., 2022; Ram et al., 2017) and environmental stability (De Frenne et al., 2019) of wooded habitats, which appear to buffer bird populations against demographic fluctuations much more effectively than heavily managed arable lands. Ultimately, the landscape heterogeneity is a key factor in driving favourable population trends, and its universal role in maintaining biodiversity provides an important conservation principle for human-modified ecosystems (Stein et al., 2014).

By contrast, urbanised areas and landscapes in which intensive agriculture predominates over forests, constrained community-wide population growth. These simplified environments are characterised by low structural complexity and heightened vulnerability to intense anthropogenic stressors. In intensive agricultural systems, widespread practices such as land drainage, heavy mechanisation, and the extensive use of pesticides directly reduce the availability of invertebrate prey, thereby lowering breeding success (Møller et al., 2021; Monnet et al., 2026; Rigal et al., 2023). Similarly, urbanisation has a negative efiect on most bird species and leads to homogenisation of bird communities (Sidemo-Holm et al., 2022), while also exposing them to severe physiological and reproductive stressors. The replacement of natural vegetation with impermeable surfaces eliminates critical foraging and nesting habitats, drastically degrading the fundamental carrying capacity for most of the avian community (Aronson et al., 2014; Corsini & Szulkin, 2025). Additional urban-associated stressors, such as artificial light at night (ALAN), pollution, and anthropogenic noise, can adversely affect avian physiology, reproductive success, and survival (De Jong et al., 2017; Dominoni et al., 2020; Gaston et al., 2017; Halfwerk et al., 2011; Isaksson, 2015).

In addition to landscape structure, persistently warmer and more humid spatial baselines drove more favourable population trends. In temperate regions, this positive demographic response is largely mediated by the species-energy relationship. Warmer and wetter environments support higher net primary productivity, consequently yielding greater biomass of invertebrate prey (Evans et al., 2005). Additionally, higher temperatures can significantly reduce overwinter mortality for resident species, while also advancing spring phenology (Lehikoinen et al., 2016). This thermal shift extends the reproductive window, enabling multi-brooded species to increase their annual fecundity (Halupka et al., 2023). However, the climate change affect bird communities unevenly, with its impact depending on species thermal niches and functional traits (Davey et al., 2012; Pearce-Higgins et al., 2015; Stephens et al., 2016). Many avian communities experience significant changes in species composition, including declines of cold-dwellers, northward or upslope shifts in their breeding ranges, alongside increases in numbers of thermophilic species (Alba & Chamberlain, 2025; Cours et al., 2025). Majority of long-distance migrants are declining (Jørgensen et al., 2016; Lemoine et al., 2007), which might be a result of ecological mismatches caused by a lack of synchronisation between peaks in food abundance and their time of arrival (Saino et al., 2011). Moreover, most birds are unable to keep up with the rapid pace of climate change (Cohen & Jetz, 2025). Together, these mechanisms demonstrate that, while climatic conditions act as general macroecological filters, they also actively reshape community composition.

### Short-term perturbations and community stability

Although spatial baselines determine the overall carrying capacity, temporal environmental deviations drive short-term population fluctuations. Our analysis revealed that temperature anomalies and transitions in the forest-cropland gradient had a negligible effect at the community level. This apparent stability does not imply an absence of demographic response. Rather, it reflects a zero-sum dynamic where divergent species-specific responses effectively cancel each other out. Since species occupy distinct ecological niches, any temporal deviation that benefits one species will inevitably disadvantage another. This dynamic is a manifestation of the temporal storage effect (Chesson, 2000), in which environmental fluctuations buffer the community and promote coexistence by shifting the competitive advantage from year to year. Consequently, the regional avian community exhibits pronounced compensatory dynamics (Gonzalez & Loreau, 2009), functioning much like an ecological portfolio (Schindler et al., 2015), effectively masking interannual temperature variability and gradual successional shifts.

Conversely, temporal changes in urbanisation and environmental heterogeneity reduced community-wide growth rates. The negative impact of urban expansion is intuitive, as rapid urbanisation-related perturbations generate anthropogenic stressors that can directly drive population declines. However, the shifts in environmental heterogeneity may result from both natural disturbances (e.g. succession, tree windbreaks, fires) and anthropogenic causes, such as local changes in land-use or forest management. Our findings demonstrate that these short-term disturbances, which increase landscape diversity, can have negative consequences for bird populations, and this effect is likely due to disruptions to local environmental optima. Many species have unimodal preferences for landscape complexity, so any sudden change in heterogeneity moves them away from the optimum, which result in a community-wide demographic cost.

At the community level, precipitation anomalies acted as a negative stressor. This negative meta-effect is driven by asymmetric species vulnerability. While the majority of the community is relatively unaffected, a subset of highly sensitive species experiences pronounced demographic declines during exceptionally wet years. High rainfall during the nesting period imposes severe thermoregulatory stress on nestlings and restricts the foraging efficiency of specific functional guilds, drastically reducing their annual recruitment (Silber & Boyle, 2026).

These findings provide an important principle for conservation management: the long-term composition of a landscape has a greater impact on bird populations than short-term disturbances. Therefore, spatial planning that prioritises the preservation and restoration of structurally complex and heterogeneous landscapes may provide particularly substantial long-term conservation benefits. Policymakers must also recognise that, although phenomena such as temperature anomalies and large-scale afforestation have limited effects at the community level, such apparent stability may obscure both underlying species turnover and a decline of highly vulnerable species. These patterns underscore the importance of continuous species-specific monitoring to evaluate the population-level impacts of environmental drivers.

### Limitations

Our analytical framework uses a massive spatiotemporal dataset to decouple carrying capacity from environmental perturbations. However, this approach is not without certain limitations. Working at 1-km spatial resolution hinders the detection of microecological patterns, such as those generated by forest age structure or changes in plant species composition, such as crop rotation. Similarly, disturbances occurring at fine temporal scales may remain undetected, despite potentially having impact on local populations. Furthermore, our environmental covariates might do not encompass a full set of potential factors influencing bird populations, for instance we do not capture localised, agricultural stressors like fertiliser loading or pesticide use, which are known to severely reduce densities of farmland birds (Boatman et al., 2004; Monnet et al., 2026). In addition, our species-specific demographic models do not explicitly parameterise biotic interactions. Finally, the robust error structures required for the statistical modelling restrict the analysis to common species. Although widespread species are responsible for most regional compensatory dynamics and ecosystem functioning (Gaston, 2010), this restriction means that our meta-analytic estimates likely represent a conservative assessment. As demonstrated by the asymmetric species vulnerability to precipitation or temperature, highly specialised species are presumably even more susceptible to the perturbations identified here.

### Policy implications

Although the landscape in Poland and broader Central and Eastern Europe remains relatively heterogeneous compared to Western Europe, rapid agricultural intensification, land consolidation, and monoculture expansion are driving severe landscape simplification. As our results demonstrate, the degradation of the baseline habitat heterogeneity limits the carrying capacity of the avian community. Therefore, in regions undergoing socio-economic and agricultural transitions, the main objective of conservation measures should be protection rather than restoration. The Common Agricultural Policy (CAP) and regional agri-environmental programmes must prioritise the maintenance of already existing habitat mosaics, ecotones, and buffer zones. In regions that have already been degraded, efforts should be made to increase landscape diversity through appropriate land-use planning.

Furthermore, urbanisation was identified as the only environmental driver with a consistently negative demographic impact on birds, acting both as a long-term spatial constrain and a temporal stressor. Suburban sprawl is accelerating rapidly across Eastern Europe, and this trend is projected to continue in the following decades (Zhou et al., 2019). Although it is difficult to reverse these suburbanisation trajectories, spatial planning can mitigate their negative consequences on avifauna by deliberately integrating heterogeneity into expanding urban areas. This can be achieved by inclusion of green-blue infrastructure, such as interconnected urban parks, rain gardens and ecological corridors. These features can provide the structural complexity needed to counteract severe carrying capacity limitations imposed by the urban development (Alba et al., 2025; Guo et al., 2025). Ultimately, safeguarding long-term regional biodiversity requires policies that recognises spatial heterogeneity not as a by-product of historical land use, but as a critical and valuable ecological asset that must be protected.

## Acknowledgements

The study was supported by the National Science Centre (NCN) in Poland (grant no. 2018/29/B/NZ8/00066). The computational resources used in this work were provided by the Poznań Supercomputing and Networking Centre (grant no. pl0090-01). The authors gratefully acknowledge MPPL volunteers for collecting the data, Chief Inspectorate for Environmental Protection (GIOŚ) and OTOP BirdLife Poland for providing the bird data.

## Author contribution statement

**Katarzyna Malinowska**: Conceptualization, Data curation, Formal analysis, Methodology, Software, Visualisation, Writing – original draft. **Tomasz Chodkiewicz:** Data curation, Validation, Writing – original draft. **Lechosław Kuczyński:** Conceptualization, Data curation, Formal analysis, Funding acquisition, Methodology, Software, Supervision, Writing – original draft

## AI use statement

Artificial intelligence tools (DeepL, ChatGPT and Gemini) were used as a language assistance tools to improve clarity and readability of the manuscript. All scientific content, including analyses and conclusions were developed and verified by the authors, who take full responsibility of the manuscript.

## Declaration of competing interest

The authors declare no conflict of interest.

## SUPPLEMENTARY MATERIAL

## Bird data

A total number of 84 common Western Palearctic bird species were used in the study, mainly passerine birds (see Table S1 for a detailed list of species). Data on bird population originate from the Common Breeding Bird Survey in Poland (MPPL), covering 25 years of monitoring (2020-2024). MPPL follows the methodology of the Pan-European Common Bird Monitoring Scheme (PECBMS; Brlík et al., 2021). In Poland, the programme has been conducted annually since 2000 and it is coordinated by OTOP BirdLife Poland on behalf of the Chief Inspectorate for Environmental Protection (GIOŚ). The monitoring scheme is based on bird counting during the breeding season along two transects located within 1 km^2^ survey plots selected by stratified random sampling. Ornithologists visit the plots twice, first between 10 April and 15 May and second between 16 May and 30 June to note all the bird species they have seen or heard, along with the distance band at which the bird was observed.

**Table S1.** A full list of bird species used in the study.

| No. | code | EURING code | English name | Latin name | mean abundance | mean prevalence |
| --- | --- | --- | --- | --- | --- | --- |
| 1 | CCC | 1340 | White Stork | *Ciconia ciconia* | 0.64 | 0.29 |
| 2 | PX | 3670 | Grey Partridge | *Perdix perdix* | 0.22 | 0.10 |
| 3 | CR | 3700 | Common Quail | *Coturnix coturnix* | 0.33 | 0.19 |
| 4 | PF | 3940 | Common Pheasant | *Phasianus colchicus* | 1.25 | 0.45 |
| 5 | GR | 4330 | Common Crane | *Grus grus* | 1.35 | 0.35 |
| 6 | CO | 6680 | Stock Dove | *Columba oenas* | 0.24 | 0.11 |
| 7 | CP | 6700 | Common Woodpigeon | *Columba palumbus* | 5.05 | 0.89 |
| 8 | SD | 6840 | Eurasian Collared-dove | *Streptopelia decaocto* | 1.89 | 0.45 |
| 9 | ST | 6870 | European Turtle-dove | *Streptopelia turtur* | 0.12 | 0.08 |
| 10 | CU | 7240 | Common Cuckoo | *Cuculus canorus* | 1.43 | 0.66 |
| 11 | AA | 7950 | Common Swift | *Apus apus* | 2.10 | 0.24 |
| 12 | U | 8460 | Common Hoopoe | *Upupa epops* | 0.28 | 0.18 |
| 13 | J | 8480 | Eurasian Wryneck | *Jynx torquilla* | 0.19 | 0.14 |
| 14 | PV | 8560 | Eurasian Green Woodpecker | *Picus viridis* | 0.16 | 0.13 |
| 15 | DM | 8630 | Black Woodpecker | *Dryocopus martius* | 0.37 | 0.27 |
| 16 | DA | 8760 | Great Spotted Woodpecker | *Dendrocopos major* | 1.80 | 0.62 |
| 17 | DE | 8830 | Middle Spotted Woodpecker | *Dendrocopos medius* | 0.06 | 0.04 |
| 18 | DI | 8870 | Lesser Spotted Woodpecker | *Dendrocopos minor* | 0.06 | 0.06 |
| 19 | L | 9740 | Woodlark | *Lullula arborea* | 0.70 | 0.31 |
| 20 | A | 9760 | Eurasian Skylark | *Alauda arvensis* | 14.28 | 0.79 |
| 21 | H | 9920 | Barn Swallow | *Hirundo rustica* | 8.00 | 0.75 |
| 22 | D | 10010 | Northern House Martin | *Delichon urbicum* | 2.28 | 0.31 |
| 23 | AT | 10090 | Tree Pipit | *Anthus trivialis* | 1.33 | 0.39 |
| 24 | AP | 10110 | Meadow Pipit | *Anthus pratensis* | 0.56 | 0.16 |
| 25 | MF | 10170 | Western Yellow Wagtail | *Motacilla flava* | 2.17 | 0.45 |
| 26 | MA | 10200 | White Wagtail | *Motacilla alba* | 1.15 | 0.52 |
| 27 | T | 10660 | Northern Wren | *Troglodytes troglodytes* | 1.08 | 0.38 |
| 28 | PM | 10840 | Dunnock | *Prunella modularis* | 0.19 | 0.11 |
| 29 | E | 10990 | European Robin | *Erithacus rubecula* | 2.59 | 0.57 |
| 30 | LUL | 11030 | Thrush Nightingale | *Luscinia luscinia* | 0.48 | 0.22 |
| 31 | LUM | 11040 | Common Nightingale | *Luscinia megarhynchos* | 0.36 | 0.15 |
| 32 | PO | 11210 | Black Redstart | *Phoenicurus ochruros* | 1.26 | 0.48 |
| 33 | PP | 11220 | Common Redstart | *Phoenicurus phoenicurus* | 0.57 | 0.29 |
| 34 | SR | 11370 | Whinchat | *Saxicola rubetra* | 1.42 | 0.44 |
| 35 | SQ | 11390 | Common Stonechat | *Saxicola torquatus* | 0.22 | 0.11 |
| 36 | OE | 11460 | Northern Wheatear | *Oenanthe oenanthe* | 0.14 | 0.08 |
| 37 | TM | 11870 | Eurasian Blackbird | *Turdus merula* | 4.34 | 0.86 |
| 38 | TP | 11980 | Fieldfare | *Turdus pilaris* | 2.09 | 0.43 |
| 39 | TF | 12000 | Song Thrush | *Turdus philomelos* | 2.62 | 0.70 |
| 40 | TV | 12020 | Mistle Thrush | *Turdus viscivorus* | 0.41 | 0.20 |
| 41 | HI | 12590 | Icterine Warbler | *Hippolais icterina* | 0.60 | 0.33 |
| 42 | SN | 12730 | Barred Warbler | *Sylvia nisoria* | 0.11 | 0.07 |
| 43 | SU | 12740 | Lesser Whitethroat | *Sylvia curruca* | 0.87 | 0.46 |
| 44 | SC | 12750 | Common Whitethroat | *Sylvia communis* | 2.56 | 0.68 |
| 45 | SB | 12760 | Garden Warbler | *Sylvia borin* | 0.55 | 0.28 |
| 46 | SA | 12770 | Eurasian Blackcap | *Sylvia atricapilla* | 5.41 | 0.85 |
| 47 | KS | 13080 | Wood Warbler | *Phylloscopus sibilatrix* | 1.85 | 0.45 |
| 48 | KC | 13110 | Common Chiffchaff | *Phylloscopus collybita* | 3.65 | 0.72 |
| 49 | KT | 13120 | Willow Warbler | *Phylloscopus trochilus* | 3.86 | 0.72 |
| 50 | RR | 13140 | Goldcrest | *Regulus regulus* | 0.48 | 0.19 |
| 51 | RI | 13150 | Common Firecrest | *Regulus ignicapilla* | 0.24 | 0.11 |
| 52 | M | 13350 | Spotted Flycatcher | *Muscicapa striata* | 0.30 | 0.18 |
| 53 | FH | 13490 | European Pied Flycatcher | *Ficedula hypoleuca* | 0.16 | 0.11 |
| 54 | AE | 14370 | Long-tailed Tit | *Aegithalos caudatus* | 0.20 | 0.09 |
| 55 | PL | 14400 | Marsh Tit | *Parus palustris* | 0.24 | 0.15 |
| 56 | PN | 14420 | Willow Tit | *Parus montanus* | 0.30 | 0.17 |
| 57 | PC | 14540 | Crested Tit | *Parus cristatus* | 0.44 | 0.19 |
| 58 | PA | 14610 | Coal Tit | *Parus ater* | 0.88 | 0.27 |
| 59 | PE | 14620 | Eurasian Blue Tit | *Parus caeruleus* | 1.81 | 0.60 |
| 60 | PJ | 14640 | Great Tit | *Parus major* | 4.85 | 0.88 |
| 61 | SE | 14790 | Eurasian Nuthatch | *Sitta europaea* | 0.68 | 0.32 |
| 62 | CF | 14860 | Eurasian Treecreeper | *Certhia familiaris* | 0.25 | 0.16 |
| 63 | CB | 14870 | Short-toed Treecreeper | *Certhia brachydactyla* | 0.14 | 0.09 |
| 64 | OR | 15080 | Eurasian Golden Oriole | *Oriolus oriolus* | 1.68 | 0.67 |
| 65 | LC | 15150 | Red-backed Shrike | *Lanius collurio* | 1.23 | 0.52 |
| 66 | LE | 15200 | Great Grey Shrike | *Lanius excubitor* | 0.13 | 0.10 |
| 67 | G | 15390 | Eurasian Jay | *Garrulus glandarius* | 1.62 | 0.64 |
| 68 | PIP | 15490 | Eurasian Magpie | *Pica pica* | 1.28 | 0.44 |
| 69 | COM | 15600 | Eurasian Jackdaw | *Corvus monedula* | 1.66 | 0.22 |
| 70 | COC | 15671 | Carrion Crow | *Corvus corone* | 0.78 | 0.27 |
| 71 | COX | 15720 | Common Raven | *Corvus corax* | 1.19 | 0.50 |
| 72 | S | 15820 | Common Starling | *Sturnus vulgaris* | 20.61 | 0.85 |
| 73 | PD | 15910 | House Sparrow | *Passer domesticus* | 7.37 | 0.48 |
| 74 | P | 15980 | Eurasian Tree Sparrow | *Passer montanus* | 2.69 | 0.43 |
| 75 | Z | 16360 | Common Chaffinch | *Fringilla coelebs* | 10.00 | 0.92 |
| 76 | SS | 16400 | European Serin | *Serinus serinus* | 0.64 | 0.24 |
| 77 | C | 16490 | European Greenfinch | *Carduelis chloris* | 1.23 | 0.43 |
| 78 | CC | 16530 | European Goldfinch | *Carduelis carduelis* | 1.43 | 0.44 |
| 79 | AB | 16600 | Common Linnet | *Carduelis cannabina* | 1.56 | 0.42 |
| 80 | PY | 17100 | Eurasian Bullfinch | *Pyrrhula pyrrhula* | 0.11 | 0.07 |
| 81 | CT | 17170 | Hawfinch | *Coccothraustes coccothraustes* | 0.95 | 0.38 |
| 82 | EI | 18570 | Yellowhammer | *Emberiza citrinella* | 4.96 | 0.86 |
| 83 | EH | 18660 | Ortolan Bunting | *Emberiza hortulana* | 0.35 | 0.15 |
| 84 | EC | 18820 | Corn Bunting | *Miliaria calandra* | 2.31 | 0.49 |

## Land cover data

Land cover data was obtained from the Moderate Resolution Imaging Spectroradiometer (MODIS) Land Cover Type product (MCD12Q1 Version 6.1) (Abatzoglou et al., 2018; Friedl & Sulla-Menashe, 2022). MODIS provides global annual land cover classifications at 500-m spatial resolution derived from supervised classification of MODIS Terra and Aqua reflectance data. We selected the International Geosphere-Biosphere Programme (IGBP) classification scheme (Type 1), which categorizes land surface into 17 classes, including natural vegetation, agricultural systems, and urban areas (Sulla-Menashe & Friedl, 2022).

Data granules covering the study area (tiles h18v03, h18v04, h19v03, h19v04) were acquired for the period 2001–2024 via the NASA AppEEARS platform. We processed the raw HDF files in R using the terra package (Hijmans et al., 2026). For each year, the granules were mosaicked using a modal rule to resolve overlapping pixel values and subsequently cropped to the extent of Poland.

To integrate categorical land cover information into our analytical grid, we calculated the fractional cover of each class within our sampling units. Binary masks were generated for each IGBP class and reprojected to the 1-km PUWG 1992 (EPSG 2180) reference grid using bilinear interpolation. This approach converted discrete pixel categories into continuous values ranging from 0 to 1, representing the proportional area coverage of each land cover type per grid cell.

Next, we combined data on coniferous, broadleaf and mixed forest types into a single category named “forests”. As a result, we obtained information on seven types of land cover, including open forests, transitional woodlands, grasslands and pastures, arable lands, urban areas, agri-forest mosaics and forests. The obtained variables were then subjected to sparce primary component analysis (SPCA) and further processed.

## Climatic data

Precipitation and temperature data originate from TerraClimate, which provide monthly climatic data with a spatial resolution of ∼4 km (Abatzoglou et al., 2018). Similarly to the land cover data, we imported the datasets into R using terra package (Hijmans et al., 2026), and reprojected to the Polish coordinate system (EPSG 2180) at a spatial resolution of 1km using bilinear interpolation. The mean temperatures were calculated from the minimum and maximum values and the annual average temperature was calculated for each study year (2001 - 2024). The sum of precipitation was calculated for each year, and the precipitation data were log-transformed to reduce skewness. This way we obtained annual precipitation and temperature variables that were further processed during the analysis.

**Tab S2.**
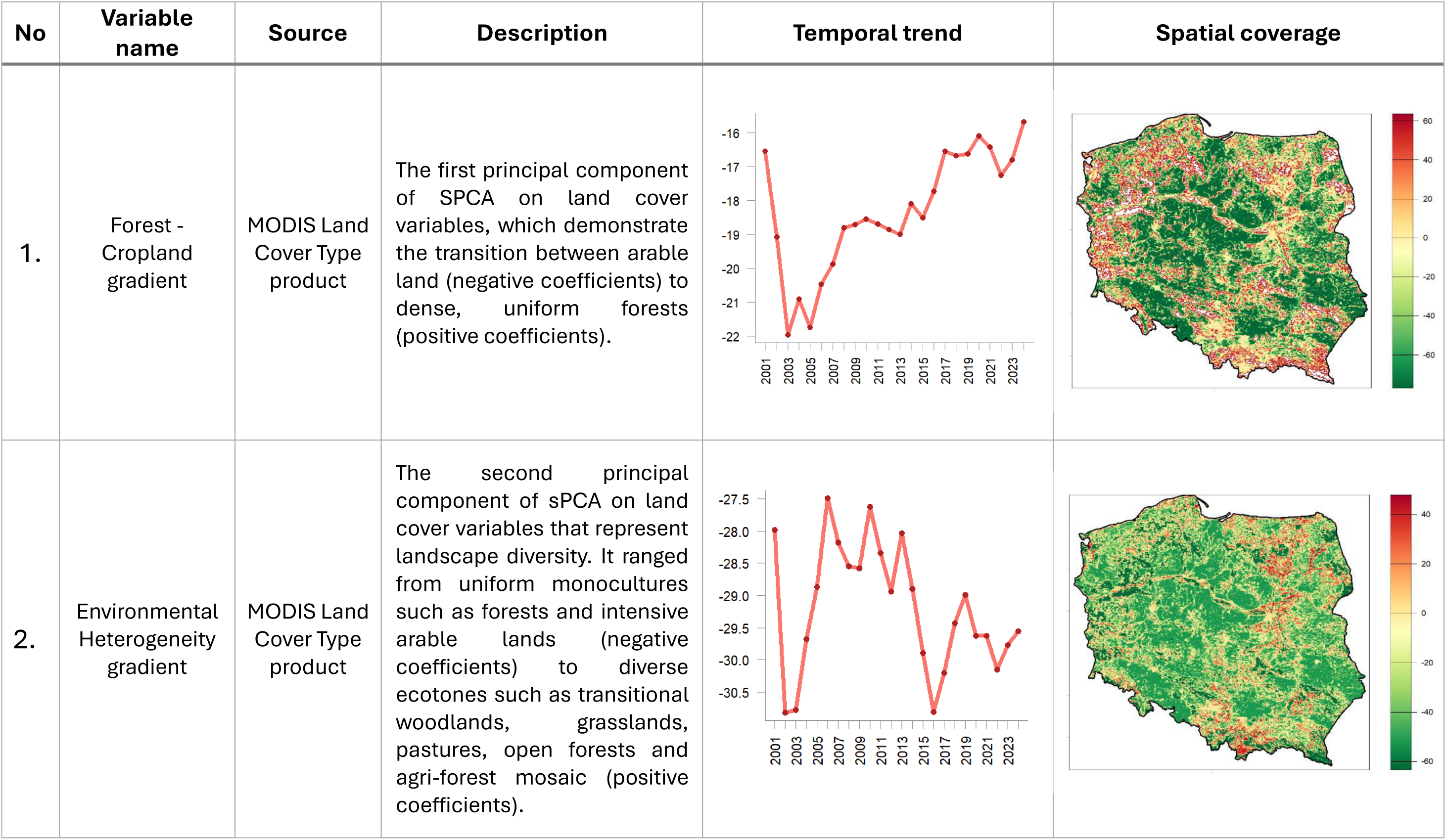
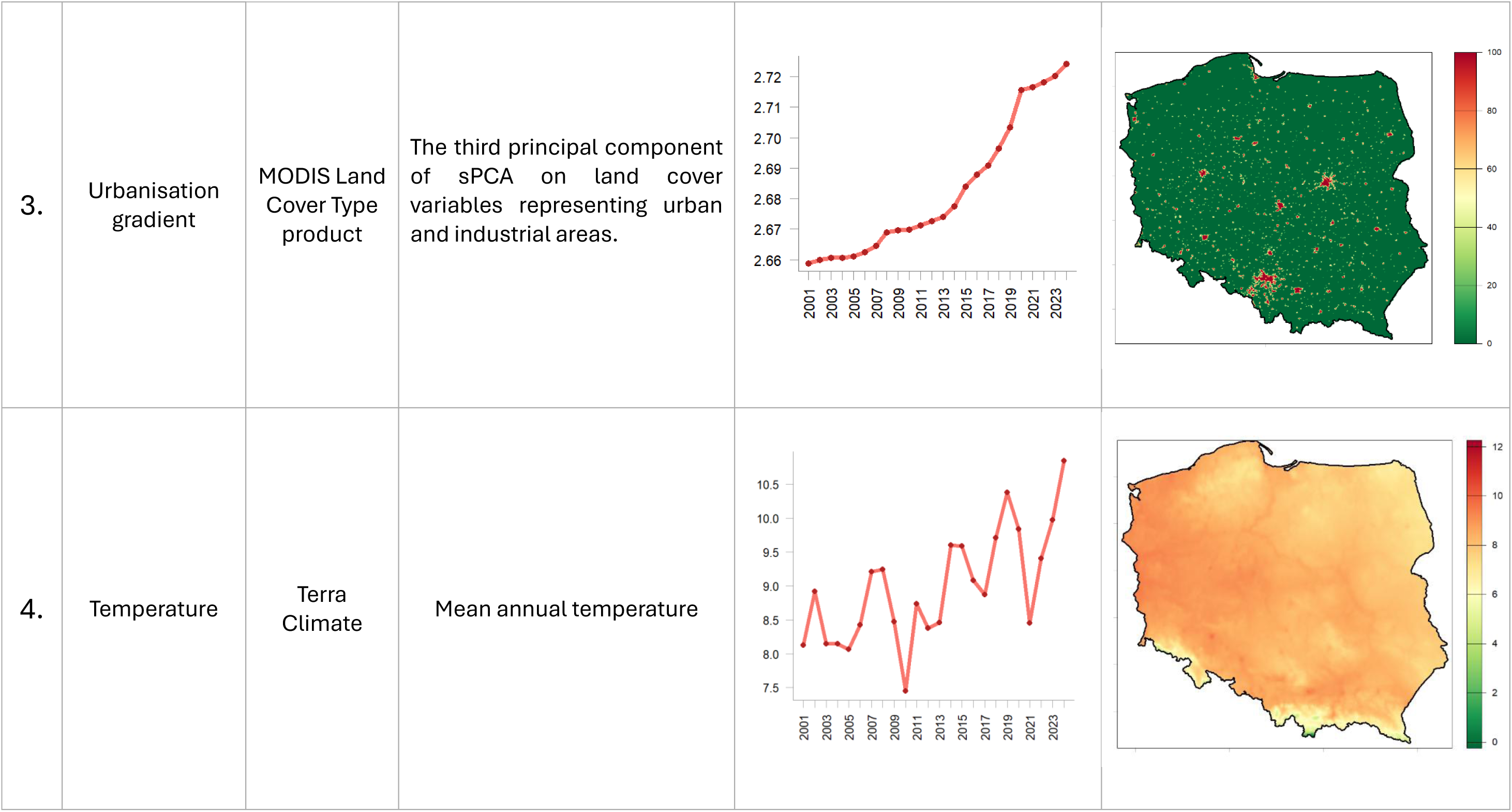
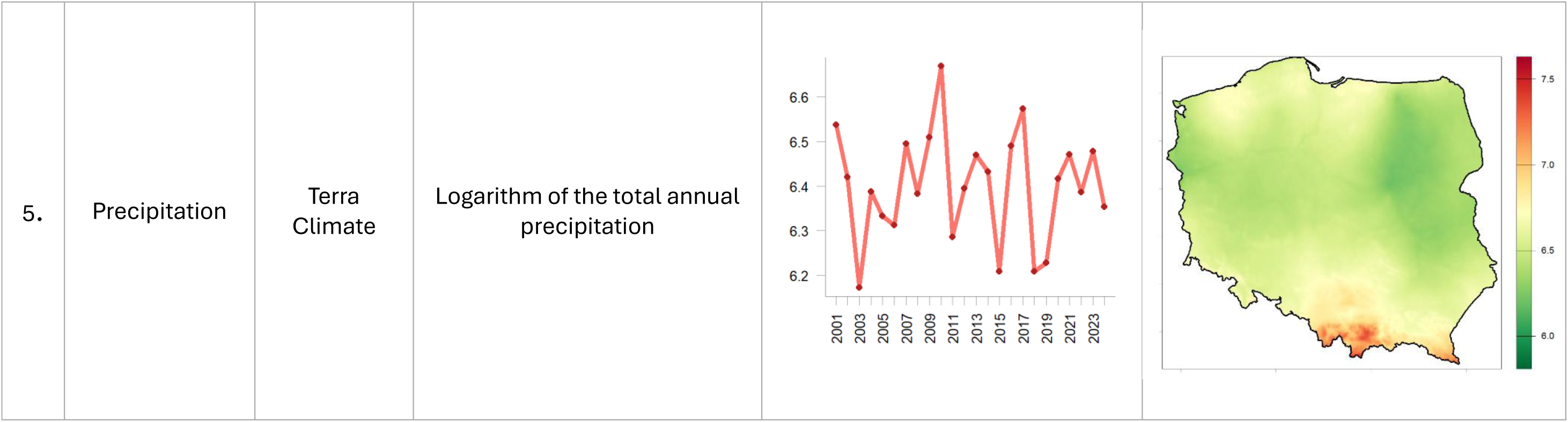
Description of five environmental drivers used in the study: PC1 (forest – cropland gradient), PC2 (environmental heterogeneity gradient), PC3 (urbanisation gradient), temperature and precipitation. Temporal trends are presented for the whole study period (2001 - 2024), whereas spatial coverage maps are shown for the midpoint of the study period, that is year 2012.

**Figure S1.**
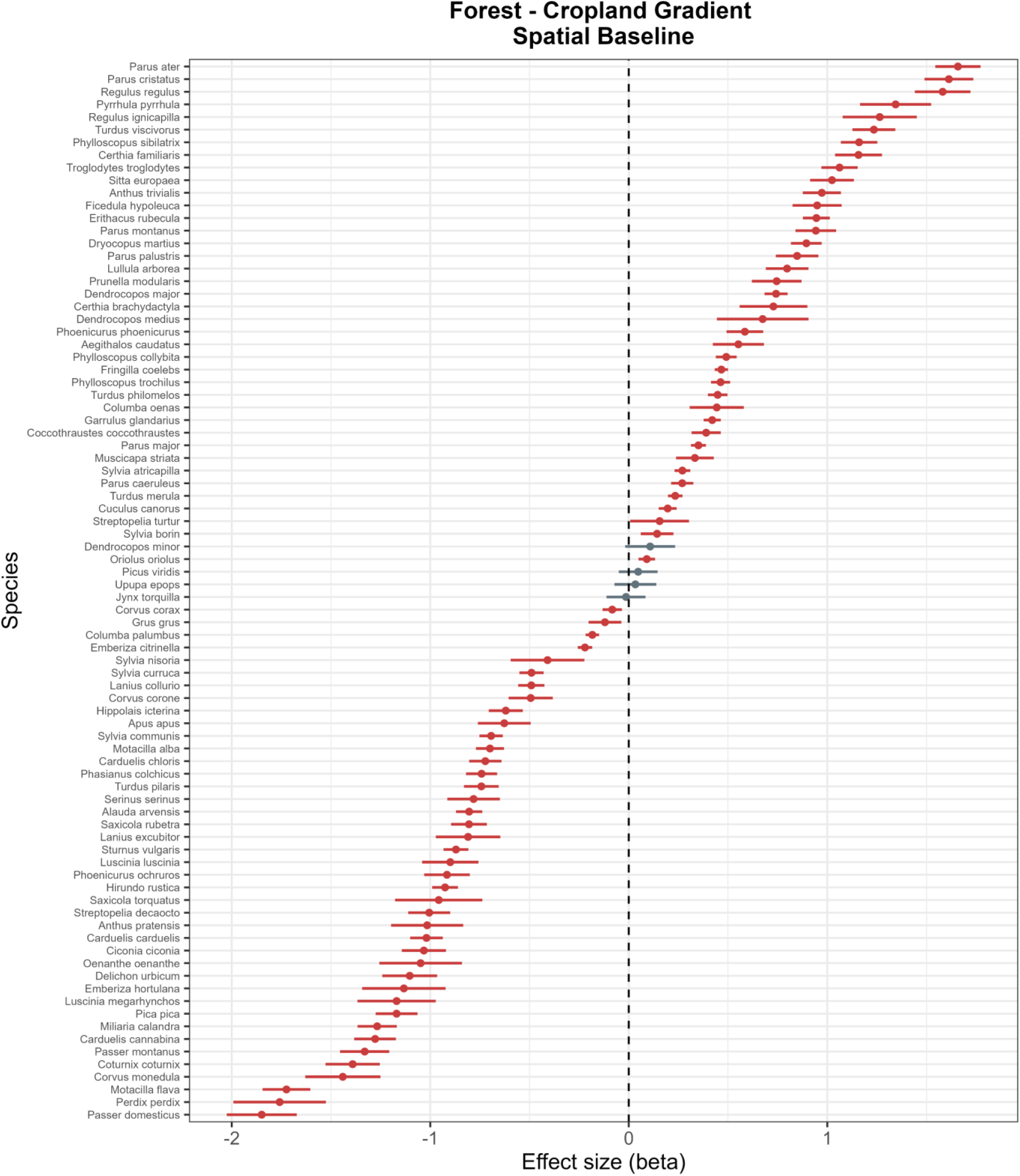
The impact of long-term spatial means of the forest – cropland gradient on bird population dynamic, estimated using species-specific demographic models. Points represent estimated effect sizes (betas), and horizontal lines indicate 95% confidence intervals. Statistically significant effects are highlighted in red, whilst non-significant effects are shaded grey.

**Figure S2.**
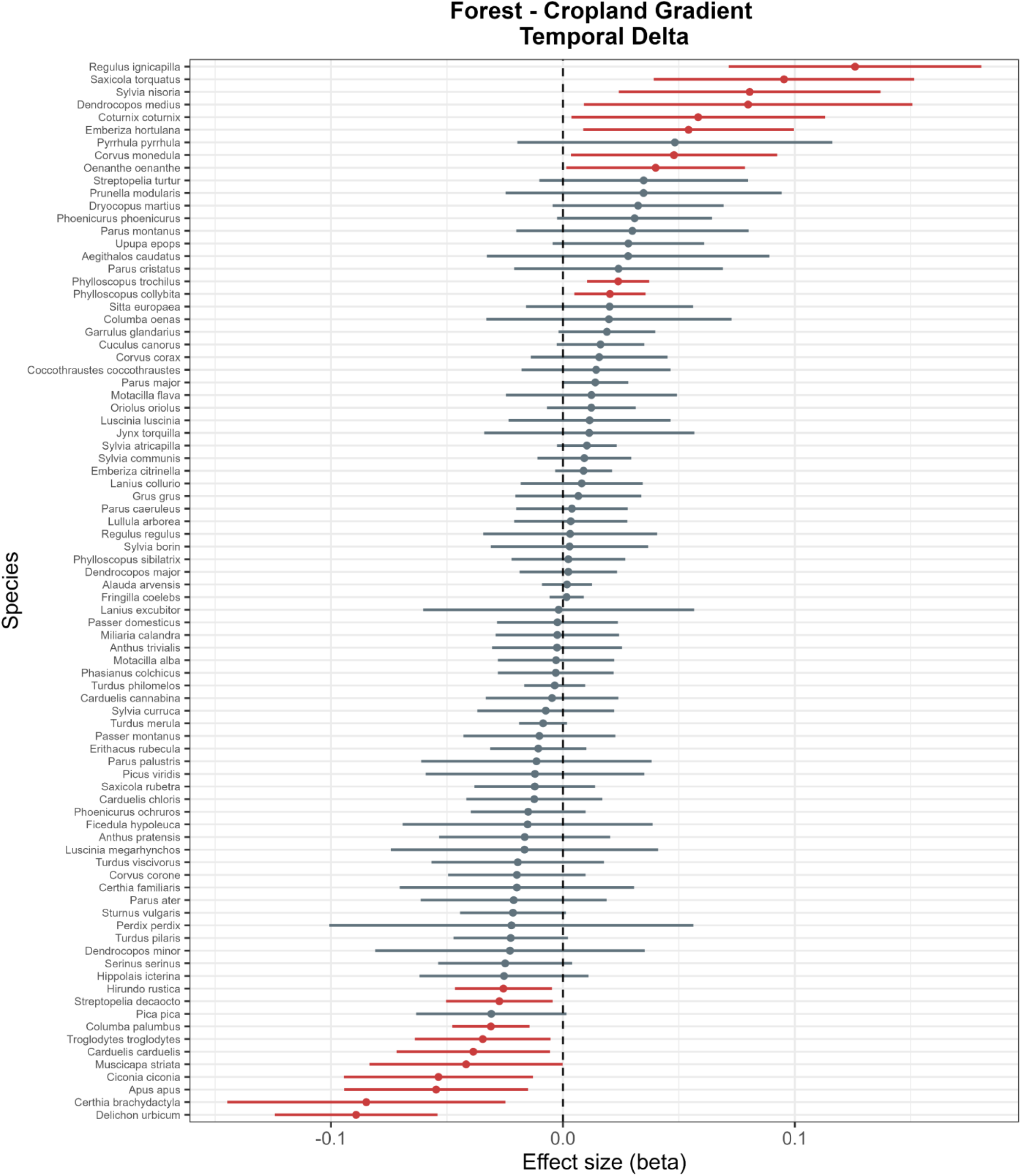
The impact of short-term disturbances in the forest – cropland gradient on bird population dynamic, estimated using species-specific demographic models. Points represent estimated effect sizes (betas), and horizontal lines indicate 95% confidence intervals. Statistically significant effects are highlighted in red, whilst non-significant effects are shaded grey.

**Figure S3.**
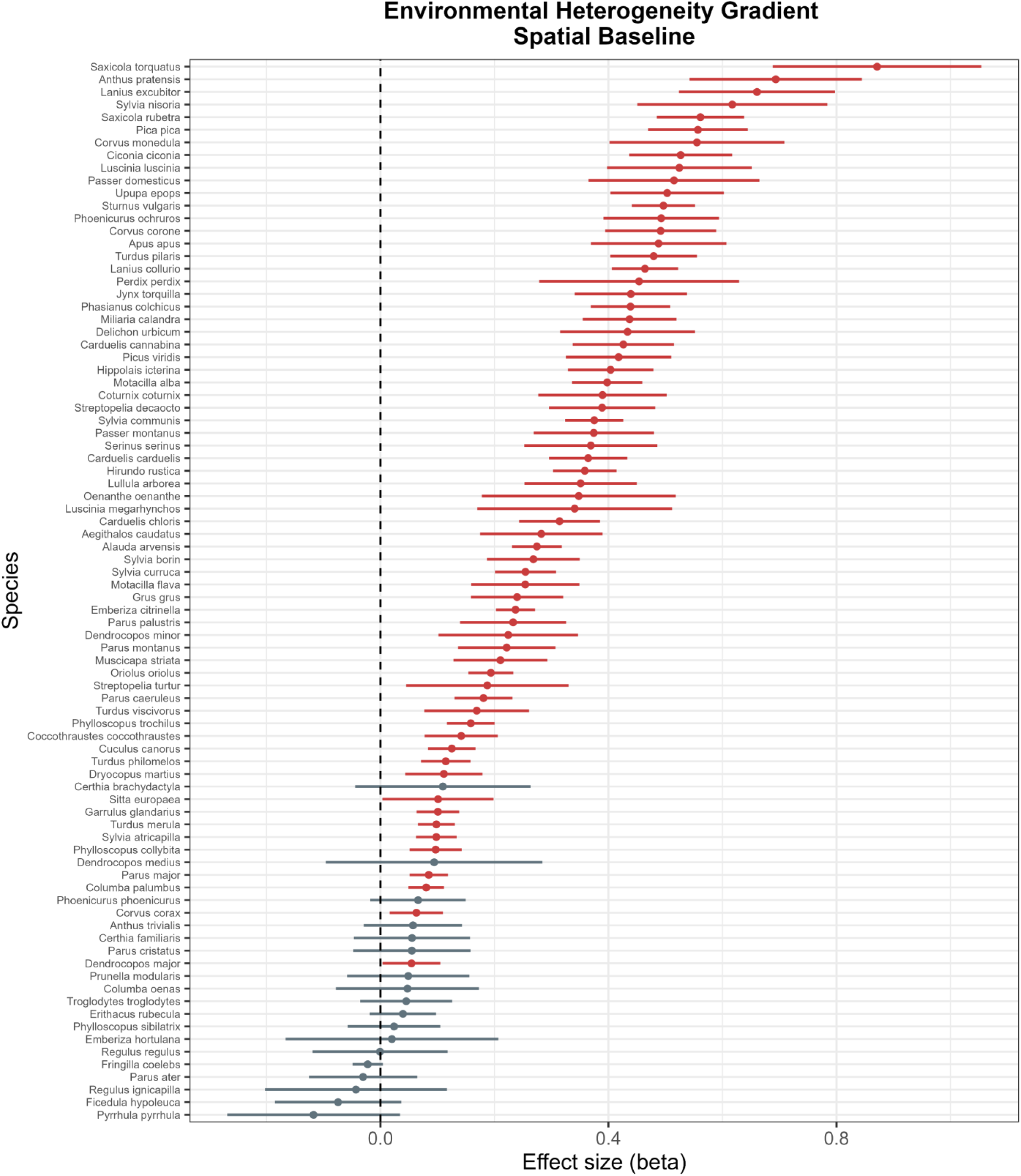
The impact of long-term spatial means of the environmental heterogeneity gradient on bird population dynamic, estimated using species-specific demographic models. Points represent estimated effect sizes (betas), and horizontal lines indicate 95% confidence intervals. Statistically significant effects are highlighted in red, whilst non-significant effects are shaded grey.

**Figure S4.**
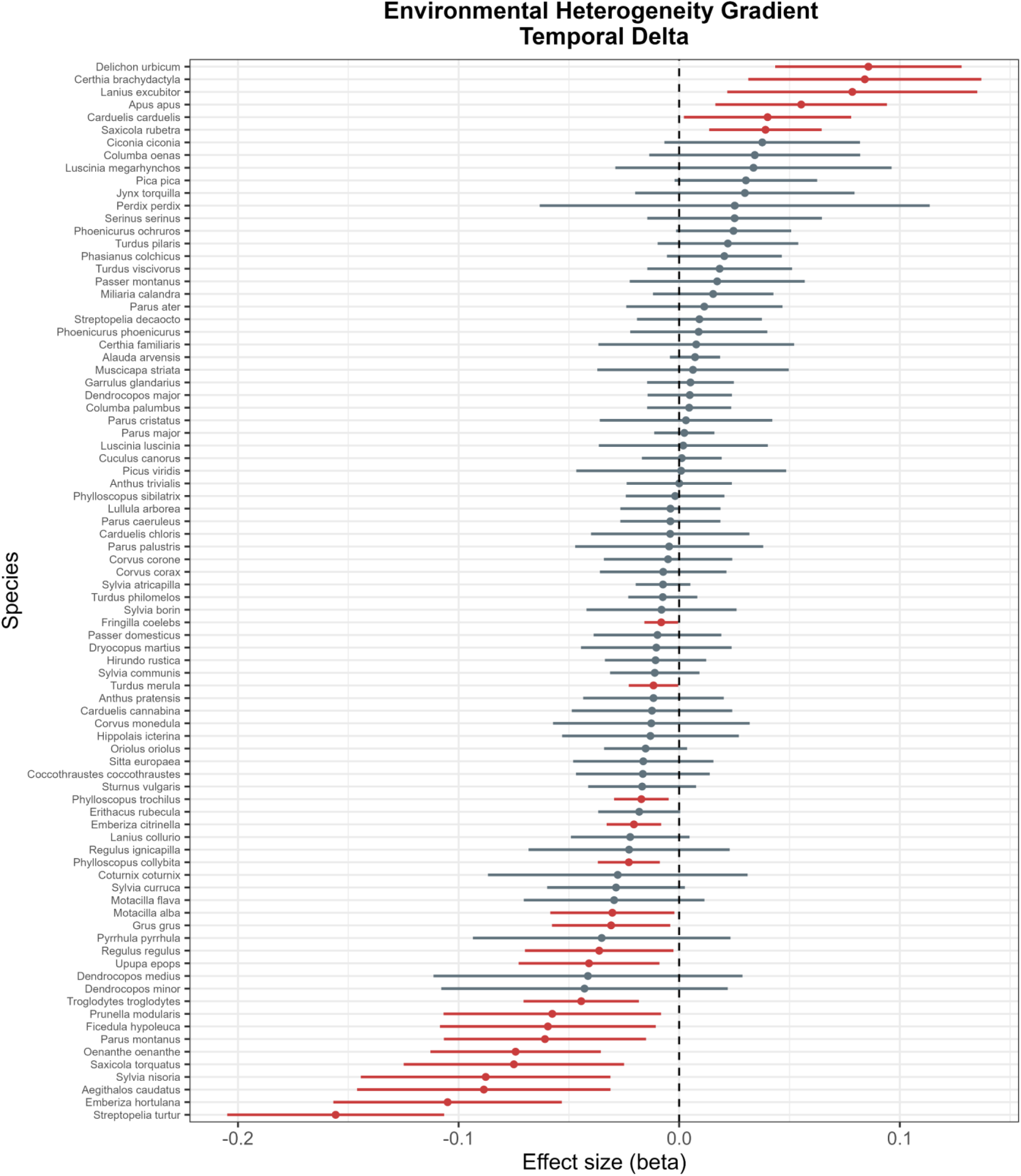
The impact of short-term disturbances in the environmental heterogeneity gradient on bird population dynamic, estimated using species-specific demographic models. Points represent estimated effect sizes (betas), and horizontal lines indicate 95% confidence intervals. Statistically significant effects are highlighted in red, whilst non-significant effects are shaded grey

**Figure S5.**
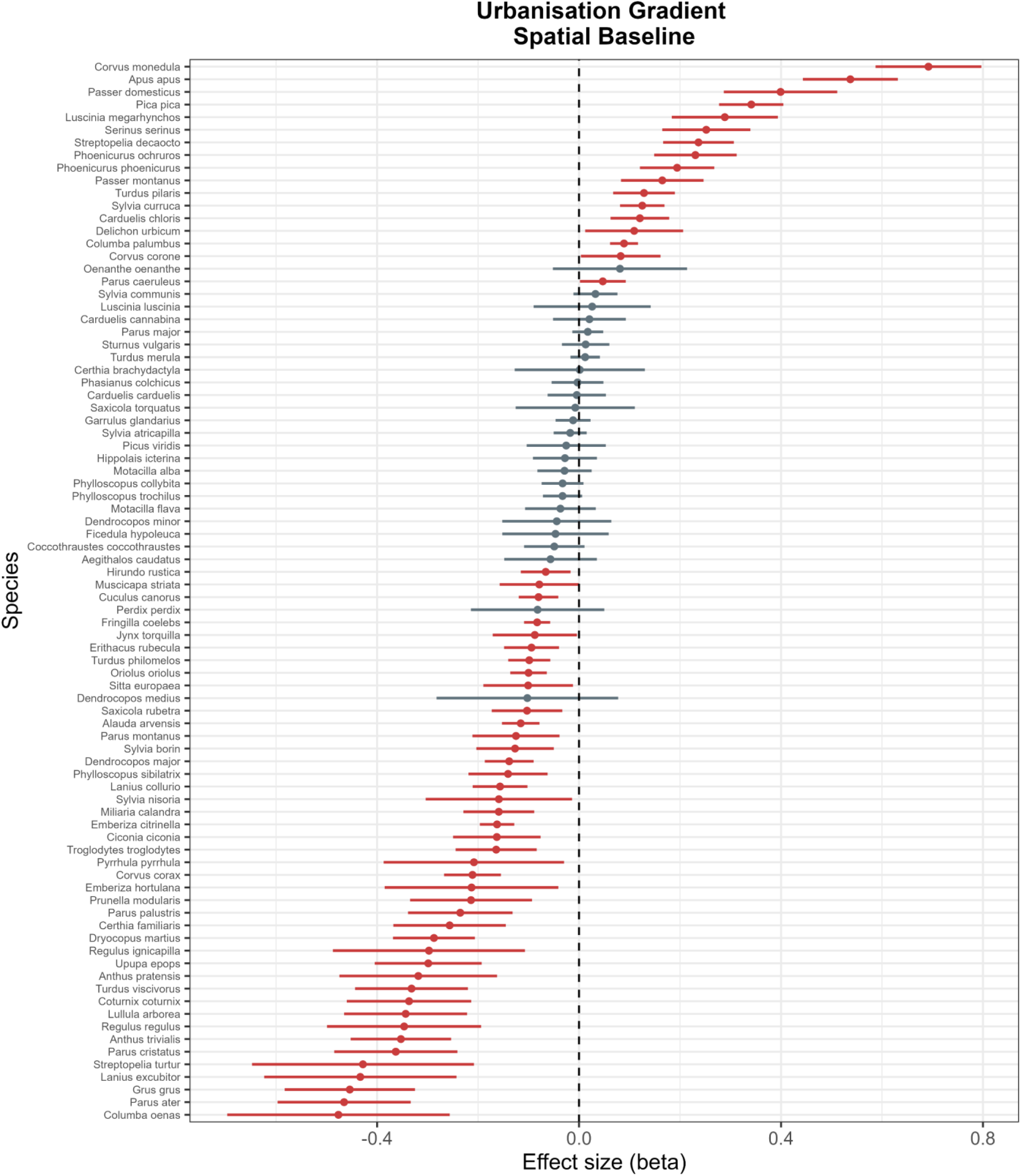
The impact of long-term spatial means of the urbanisation gradient on bird population dynamic, estimated using species-specific demographic models. Points represent estimated effect sizes (betas), and horizontal lines indicate 95% confidence intervals. Statistically significant effects are highlighted in red, whilst non-significant effects are shaded grey.

**Figure S6.**
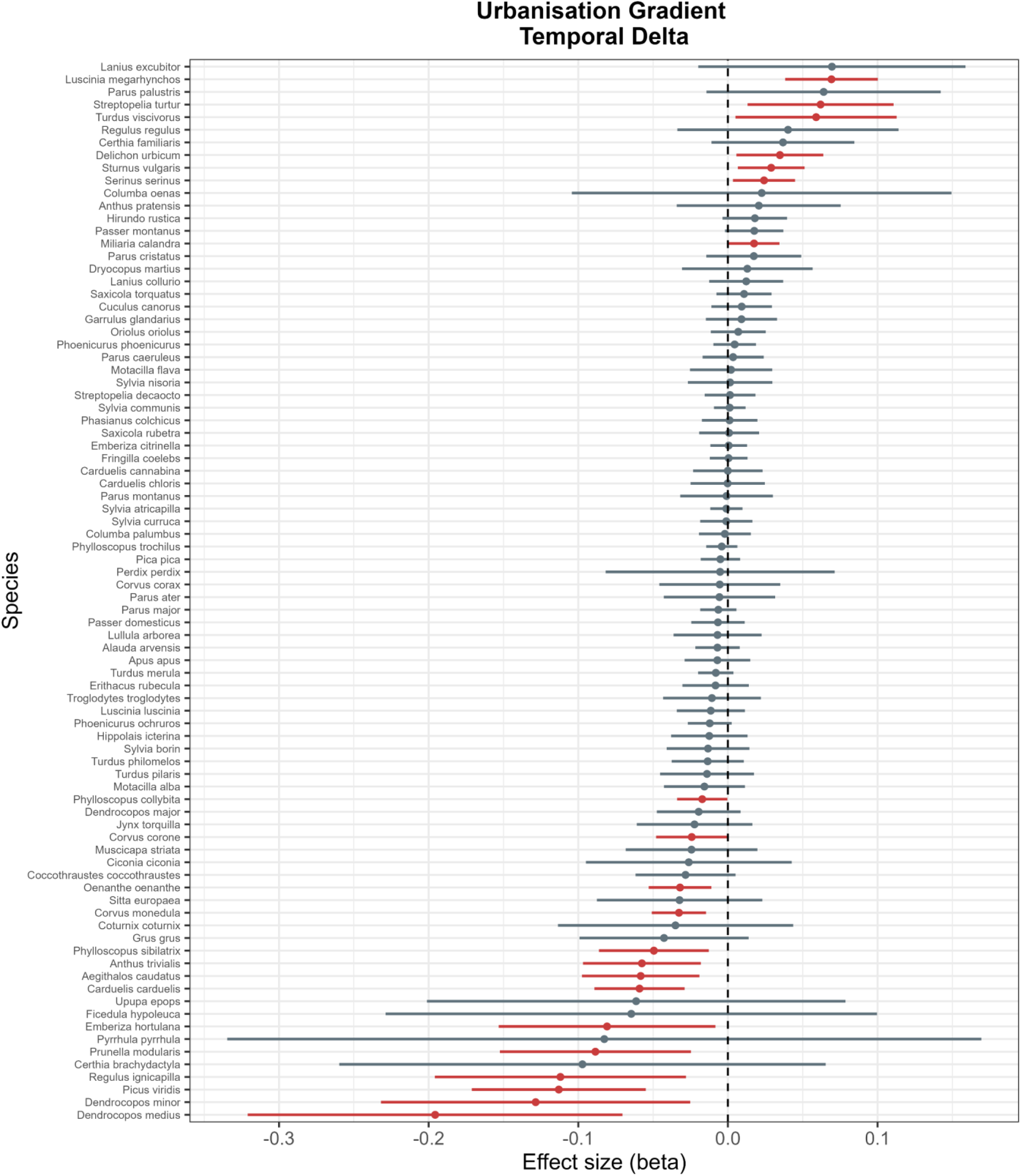
The impact of short-term disturbances in the urbanisation gradient on bird population dynamic, estimated using species-specific demographic models. Points represent estimated effect sizes (betas), and horizontal lines indicate 95% confidence intervals. Statistically significant effects are highlighted in red, whilst non-significant effects are shaded grey.

**Figure S7.**
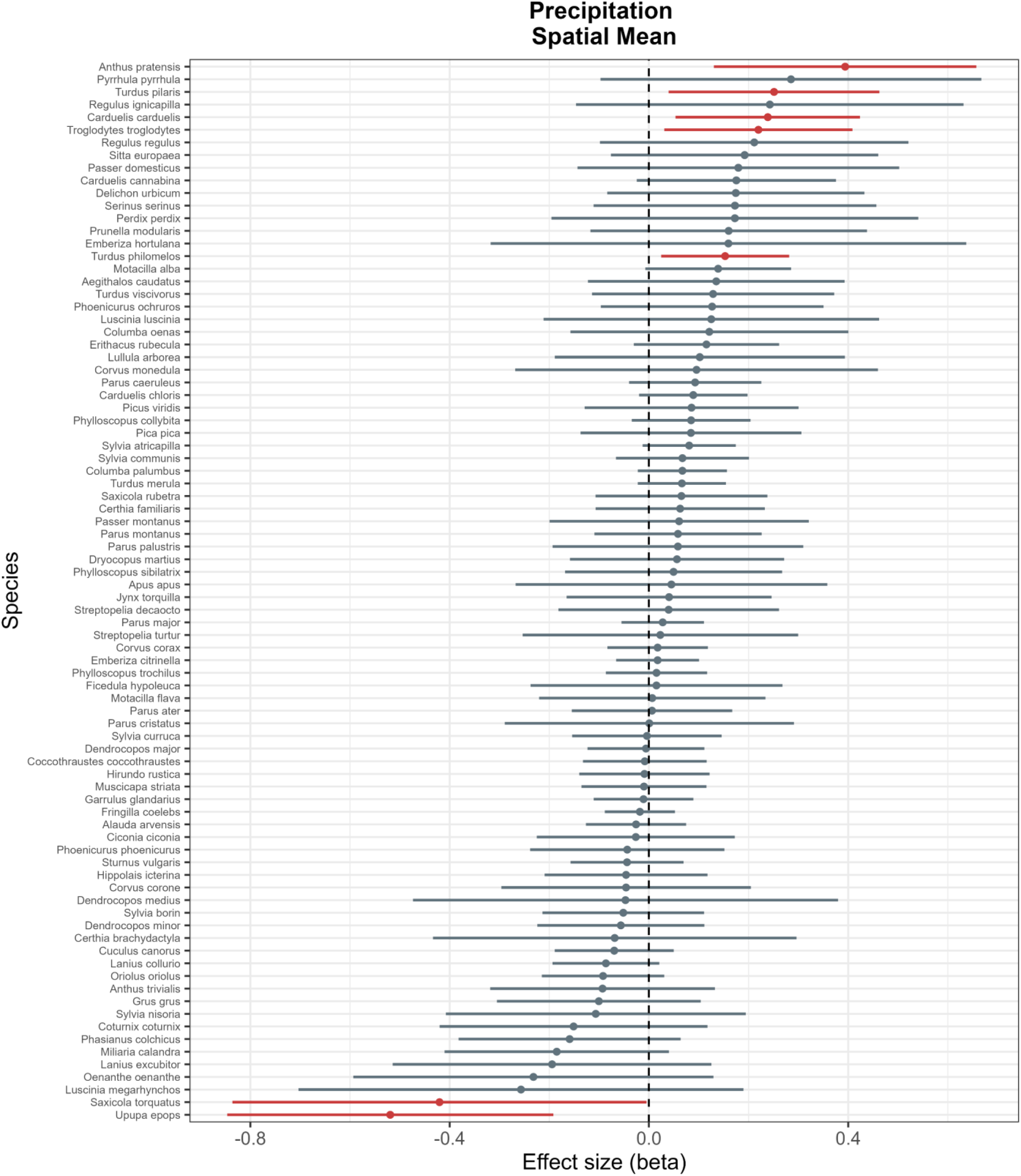
The impact of long-term spatial means of precipitation on bird population dynamic, estimated using species-specific demographic models. Points represent estimated effect sizes (betas), and horizontal lines indicate 95% confidence intervals. Statistically significant effects are highlighted in red, whilst non-significant effects are shaded grey

**Figure S8.**
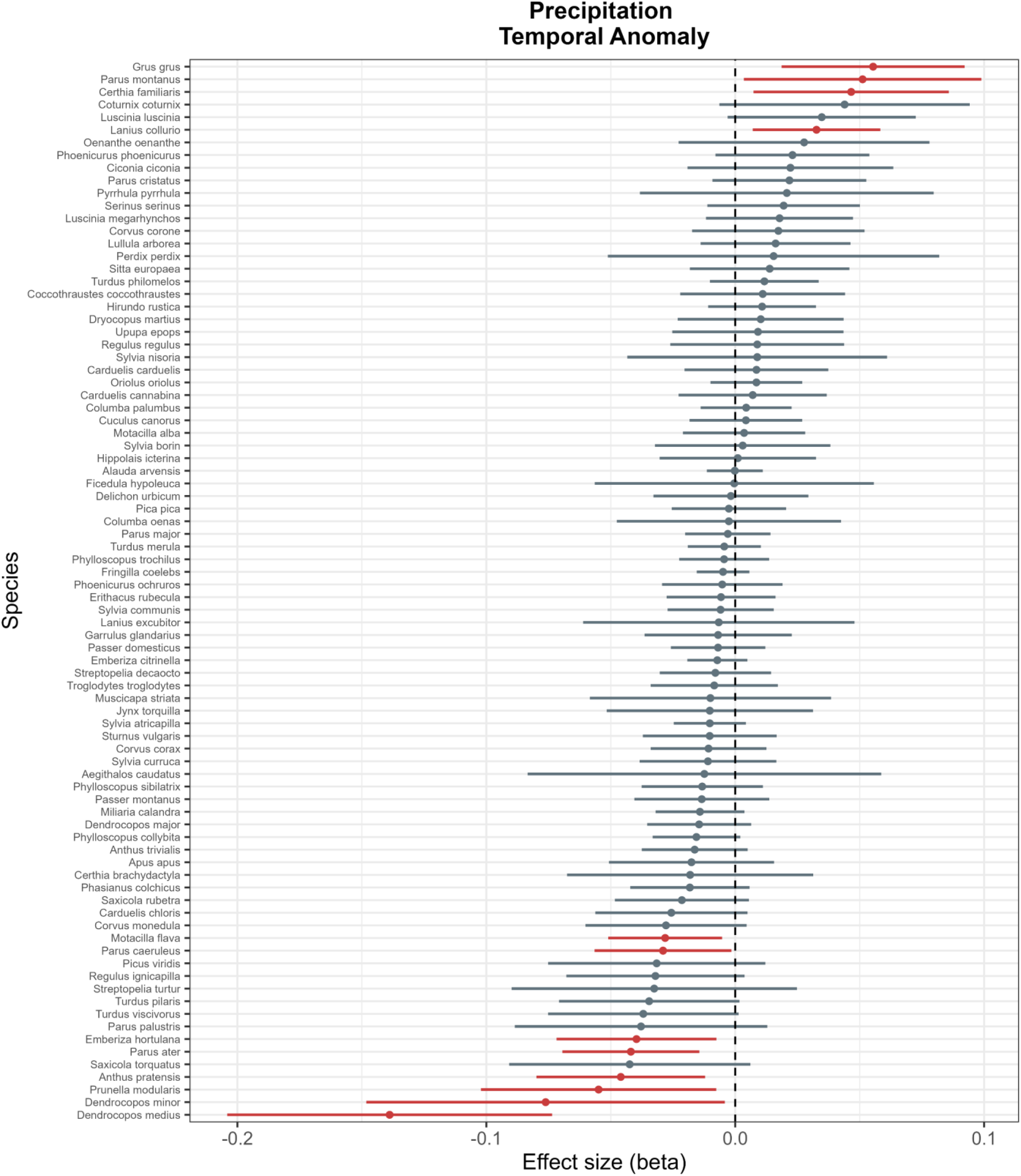
The impact of short-term precipitation fluctuations on bird population dynamic, estimated using species-specific demographic models. Points represent estimated effect sizes (betas), and horizontal lines indicate 95% confidence intervals. Statistically significant effects are highlighted in red, whilst non-significant effects are shaded grey.

**Figure S9.**
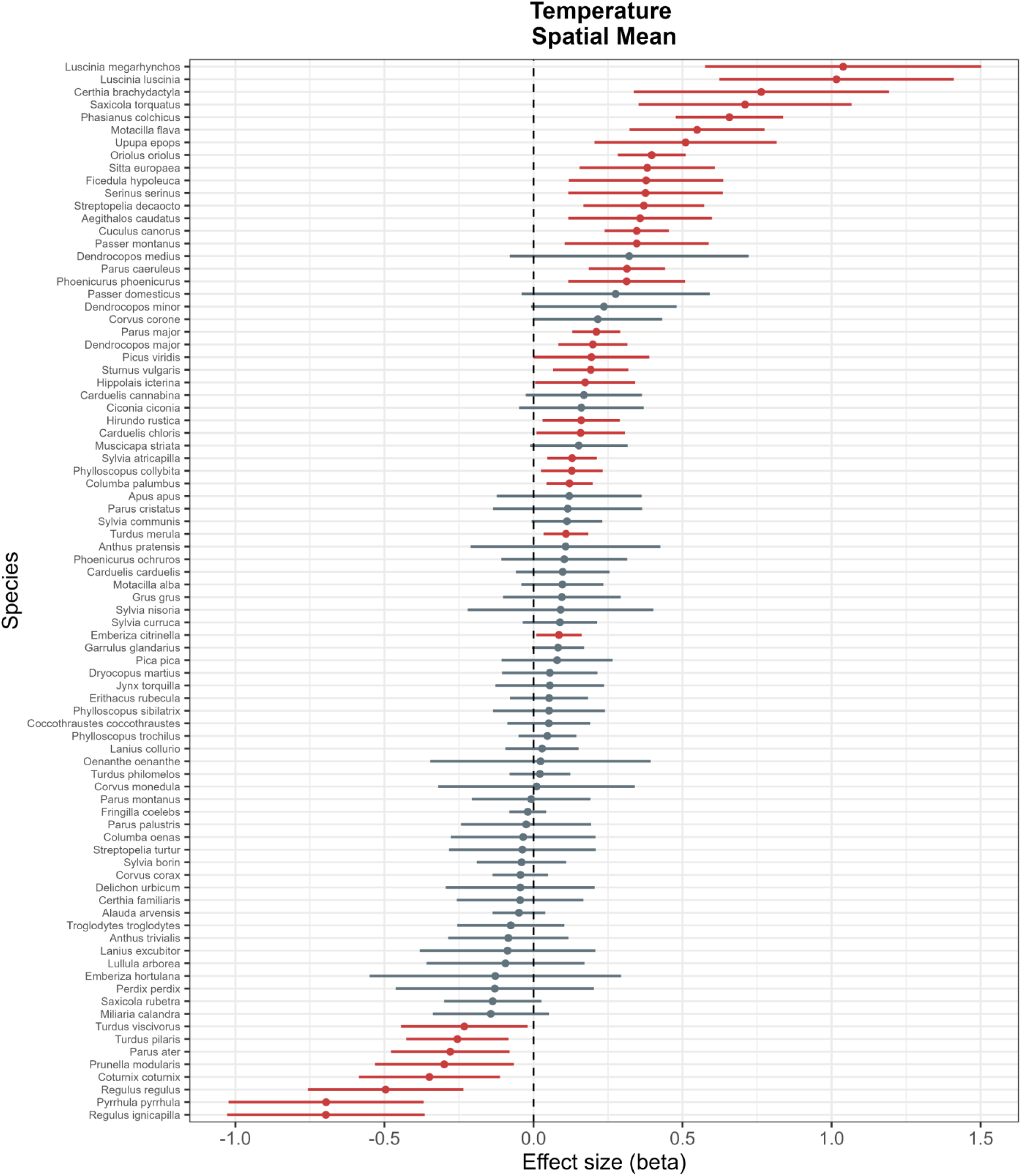
The impact of long-term spatial means of temperature on bird population dynamic, estimated using species-specific demographic models. Points represent estimated effect sizes (betas), and horizontal lines indicate 95% confidence intervals. Statistically significant effects are highlighted in red, whilst non-significant effects are shaded grey.

**Figure S10.**
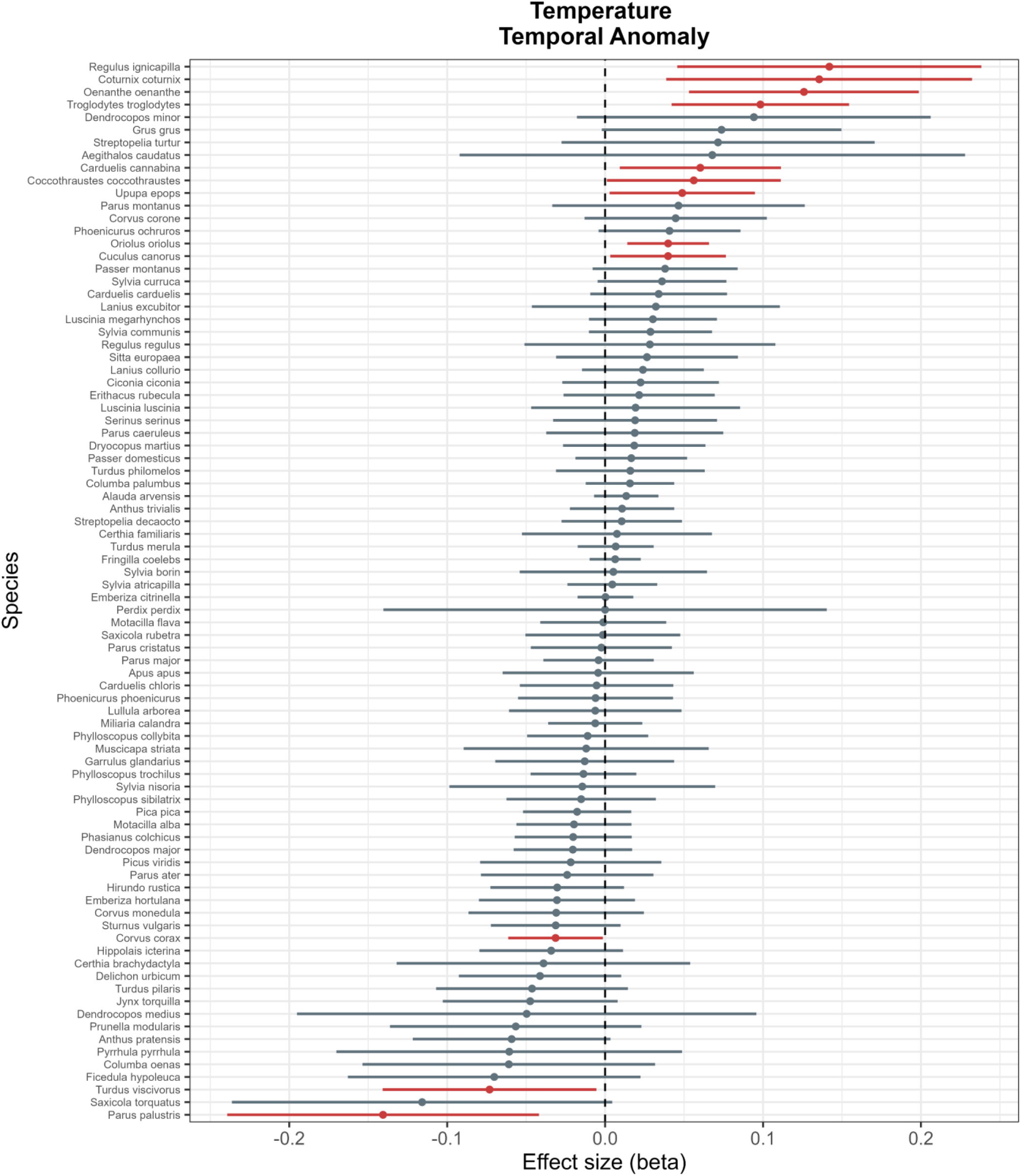
The impact of short-term temperature fluctuations on bird population dynamic, estimated using species-specific demographic models. Points represent estimated effect sizes (betas), and horizontal lines indicate 95% confidence intervals. Statistically significant effects are highlighted in red, whilst non-significant effects are shaded grey.

## Notes

### Competing Interest Statement

The authors have declared no competing interest.

